# Supra-regular recursion in primate vocal sequences

**DOI:** 10.64898/2026.09.19.752870

**Authors:** Remo Nitschke, Lothar Sebastian Krapp, Chundra Cathcart, Quentin Gallot, Alexandra B. Bosshard, Judith M. Burkart, Mélissa Berthet, Simon W. Townsend, Martin Surbeck, Klaus Zuberbühler, Balthasar Bickel

**Affiliations:** ISLE Institute, University of Zurich, Zurich & Switzerland; Department of Biology, Université de Neuchâtel, Neuchâtel & Switzerland; Department of Evolutionary Anthropology, University of Zurich, Zurich & Switzerland; Department of Philosophy, University of Milan, Milan & Italy; Department of Human Evolutionary Biology, Harvard University, Cambridge & USA

## Abstract

Complex syntax is the widely accepted demarcation between animal and human communication. Although animal vocal systems occasionally contain pockets of syntax-like structures, their grammars are thought to be at best regular, whereas human language is supra-regular, i.e., including at least context-free computational complexity. Leveraging a new likelihood method for probabilistic formal grammars, we show that the call sequences of three phylogenetically diverse nonhuman primates (bonobos, olive colobus monkeys, common marmosets) include parts that are most likely generated by supra-regular recursive grammars, similar to human language. These findings challenge syntax as the major transition in hominid evolution and opens up prospects for alternative evolutionary scenarios.

---

Supra-regular recursion has often been claimed to be unique to human language, creating a fundamental difference from the communication systems of other animals and raising challenging questions for hominin evolution (*1–5*). What is at stake is the specific kind of hierarchical structure that permeates human languages. Suppose someone tells you: “They discussed biology curricula”. It will be clear that it is curricula that are being discussed, not biology, even though *discussed* and *biology* stand right next to each other and discussing biology is an entirely plausible thing to do. The cause of this is a specific hierarchical structure: *biology* is embedded in a phrase dominated by *curricula*, and this is in turn embedded in a phrase dominated by *discussed*, yielding the nested structure [*discussed* [*biology curricula*]]. Hierarchical structures in languages have arbitrary branching directions — *curricula* follows the word it belongs to (*discussed*), while *biology* precedes the word it belongs to (*curricula*) — and they can be arbitrarily deepened by recursion, e.g. [*discussed* [[*cell biology*] *curricula*]] or even [*discussed* [[[*stem cell*] *biology*] *curricula*]].

The generation of such hierarchies requires supra-regular grammars, or what is known more technically as grammars of at least context-free complexity (*6–8*) (Box 1). While linguists use a variety of formalisms to describe human language, virtually all of them include some mechanism allowing that level of complexity in some component of their theory (*9, 10*). Context-free systems exceed regular grammars, which are limited to a single, uniform direction of branching (bracket expansion) and a fixed linear context for capturing the relationships between words or symbols. For example, a regular grammar with context window of 1 (also known as a bi-gram) could capture that *curricula* follows *biology* and a window of 2 (a tri-gram) could capture that it follows *discussed biology*, but this fails to capture the hierarchical structure that groups *discussed* with *curricula* and not *biology* over arbitrarily deeply nested insertions, as in *discussed* [^*n*^]^*n*^ *curricula*.

Previous research has demonstrated that non-human primates master supra-regular grammars when sufficiently trained on artificial visual stimuli (*11–13*). However, in line with received expectations (*1–3*), current evidence suggests that they do not generally exploit this capacity in communication. Indeed, much structure in call sequences of diverse non-human primates has been captured rather by *n*-gram models (*14–18*), which stay firmly within the regular computational capacity. Whilst this might support the notion of supra-regular communication being unique to language, progress has been hampered by a lack of methods to empirically distinguish between animal call sequences generated by a regular vs. a supra-regular grammar. In humans, native speaker judgments regarding word groupings (phrases) serve as a key litmus test for detecting supra-regular patterns in language, but such an approach is nearly impossible in other primates, due to the absence of such judgments or some other evidence for hierarchical groups of calls (e.g. in terms of usage context, meaning, or formal dependencies).

**Box 1**

Formal Language Theory as a tool for capturing animal call sequences

Formal language theory, a branch of mathematics and computer science, describes rule systems (called “grammars”) that generate sets of symbol sequences (called “languages”), usually with structure (brackets, trees) (*6*). These sets, and the grammars that generate them, show nested computational complexity: “regular” ⊊ “context-free” ⊊ “context-sensitive” ⊊ “recursively enumerable” languages (*8*). If all possible sequences of a language are known, it is possible to prove its minimal complexity (*6– 8, 19–21*). Conversely, one can design grammars that generate languages at given levels of complexity and then use these to test cognitive capacities across species (*11–13*).

In any human language, it is mostly straightforward to decide which sequences (sentences) belong to it: for example in English, we simply ask speakers to judge whether, say “they discussed curricula” or “discussed curricula they” is part of the set. But for animal call sequences we don’t know the set definition. Without this, the complexity level is demonstrably undecidable (*22*). We break out of this impasse by replacing discrete by probabilistic grammars, where each grammar rule has a nonzero probability (see *Supplementary Text S2* for the formal definition). Such a system assigns unequal probabilities to sequences (*23*).

Consider, as a minimal example, two grammars that generate hierarchically structured sequences of symbol **a** with the non-terminal (intermediate node) symbol **S**. In each case there are two rules with probabilities *p*_1_ = .6 and *p*_2_ = .4, generating for example the sequences [**a** [**a**]] and [[**a**][**a**]]. The probabilistic regular grammar (pREG) produces sequences at different lengths *n* with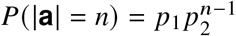, while the probabilistic context-free grammar (pCFG) produces these with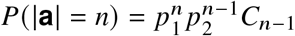, where *C* is the Catalan number (*23*). The resulting probability differences enable separation of the most likely grammar that generated a collection of sequences even if it is unknown what might define the collection as a proper set.

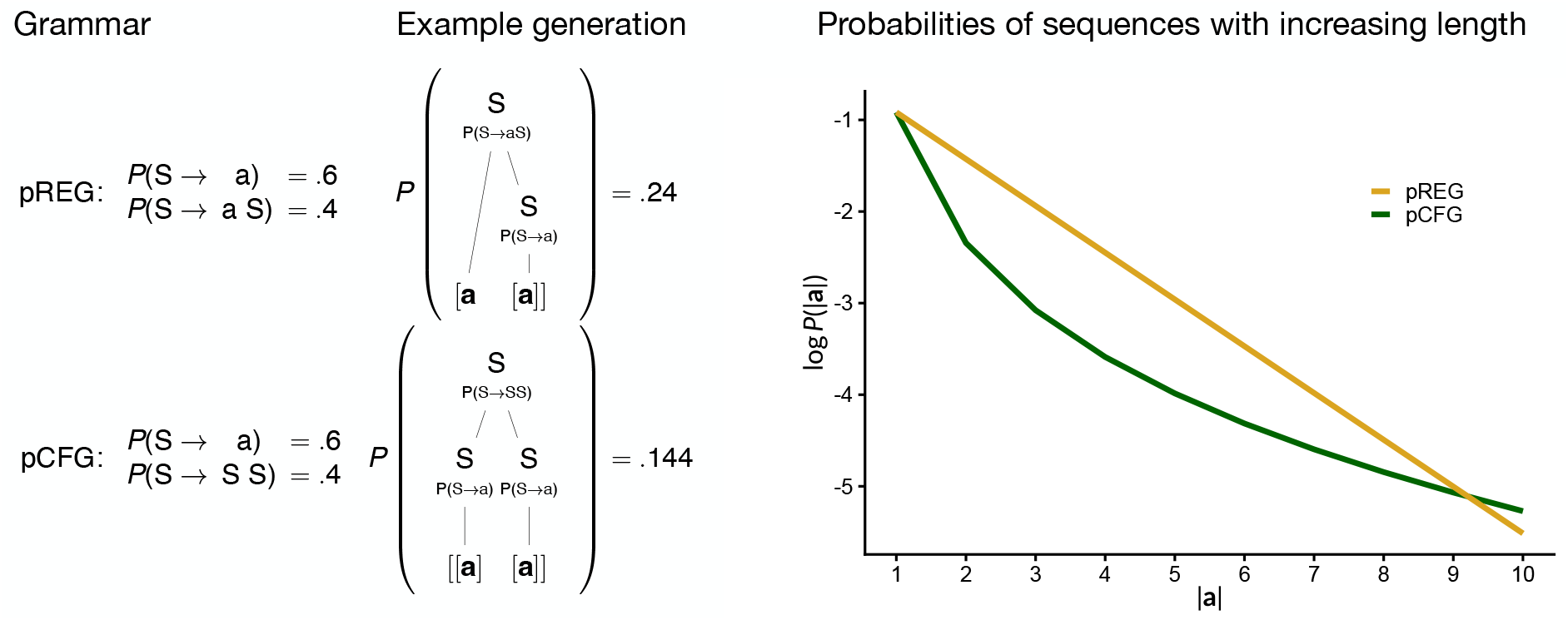

Here, we exploit recent results in probabilistic (instead of discrete) formal language research (*23*) to develop and validate a new unsupervised methodology that distinguishes sequences generated by a probabilistic regular (pREG) from sequences generated by a probabilistic context-free (pCFG) grammar (Box 1, Supplementary Text S1–S3). Critically, we apply this to naturally-produced call sequences of three phylogenetically diverse primates: bonobos (*Pan paniscus*), olive colobus monkeys (*Procolobus verus*), and common marmosets (*Callithrix jacchus*). In each species, we induce pREGs and pCFGs, maximize the log-likelihood of their parameters (rule probabilities, context window, i.e. *n*-gram size in pREG, number of non-terminal symbols in pCFG). We then compare the expected log-likelihood of each grammar over held-out data in 20-fold cross-validation, summarized as ELLD scores (expected log-likelihood differences; *Materials and Methods*). Based on validation experiments with artificial grammars (*Supplementary Text S3*) we expect a positive ELLD for the pCFG model if the data was produced by a pCFG and a negative ELLD if it was produced by a pREG.

For animal call sequences, received theory would predict that held-out sequences are more likely under pREGs than pCFG, i.e. negative ELLD scores throughout the data. Surprisingly, this is not what we found.

## Results

Figure 1A shows that the call sequences of all three species contain as many folds that are most likely generated by a pCFG as folds that are most likely generated by a pREG. The result is validated by the expected log-likelihood differences that arise when we explicitly simulate pCFGs and pREGs with a symbol inventory and sample size comparable to the primate data, taking bonobos as an example (*Supplementary Text S3* for more general validation on simulated data). This means that some parts of the data are best captured by hierarchical structures that require supra-regular mechanisms for their generation. Figure 1B illustrates some of the resulting hierarchies present in the bonobo dataset, sampled from the best-fitting pCFG for this primate.

**Figure 1.**
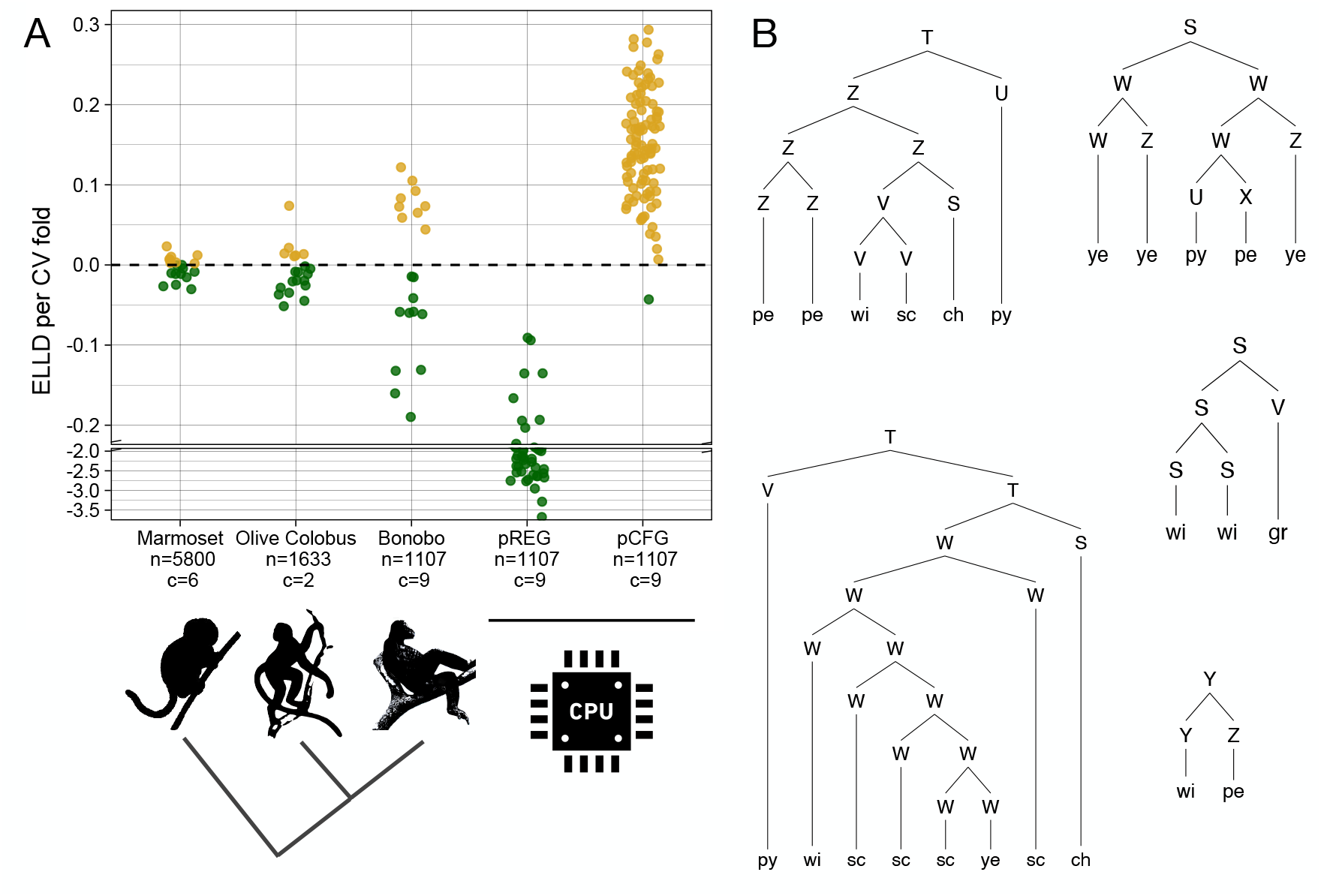
Likelihood differences between probabilistic regular (pREG) and probabilistic context-free (pCFG) grammars per cross-validation fold. **A**: For each primate there are about as many parts of the sequence data (cross-validation folds) that are most likely generated by a pCFG as by a pREG (*n*: number of sequences used for the analysis; *c*: inventory of calls; *ELLD*: expected log likelihood difference of held-out sequences in a 20-fold cross-validation; *Materials and Methods*), largely mirroring what is found when sequences are simulated from artificial grammars with the same number of calls and similar sample size as the bonobo data. **B**: A random selection of hierarchical structure in bonobo sequences (with arbitrary non-terminal symbols) sampled from the parses provided by the best-performing pCFG.

There is considerable variation in call inventory size (*c* = 2 call types in olive colobus monkeys, *c* = 6 in marmosets, *c* = 9 in bonobos), but results do not differ systematically by this. To probe this further, we estimated ELLD scores under an alternative, more fine-grained analysis of the olive colobus data, distinguishing *c* = 9 instead of *c* = 2 call types. While this slightly increased the number of folds most likely under a pREG, it didn’t change the overall pattern (*Supplementary Figure S1*). The results are also immune against differences in the number of parameters per grammar since the ELLD score is based on cross-validation, hence a direct measure of predictive performance, not model fit.

To contextualize these findings, Figure 2A shows the ELLD scores for human language data. Language does not occupy a single level of computational complexity but contains parts at very different levels (Figure 2B). Notably, sound sequences are generally assumed to be generated by a regular grammar (*24, 25*). By contrast, sequences of words or parts of speech categories (“POS” nouns, verbs, adjectives, prepositions, etc.) are assumed to be generated by at least context-free grammars (*6, 20*). To compare the complexity level of these systems with ELLD scores, we used a large database of English (*Materials and Methods*). We extracted phone sequences (transcribed discrete sound segments (e.g. jh ah s t for “just”), expecting negative ELLD scores suggesting pREGs, and POS sequences (e.g. N N V A N V), expecting positive ELLD scores suggesting pCFGs. (We did not use word sequences because unsupervised learning of grammars from these is computationally overly expensive while not allowing much insight beyond what can we can be derived more efficiently from POS sequences.) The results in Figure 2A confirm the expected differences, with phone sequences more likely generated by pREGs and POS sequences more likely by pCFGs. Intriguingly, however, when the number of sequences is down-sampled to be closer to the nonhuman primate data, parts of the POS sequence data are more likely generated by a regular grammar. Similarly, when the number of marmoset sequences is downsampled to be closer to the olive colobus and bonobo datasets, we found a decrease in the number of folds with a positive ELLD score (*Supplementary Figure S2*). This raises the possibility that larger samples of nonhuman primate sequences than currently available might show even more supra-regular parts.

**Figure 2.**
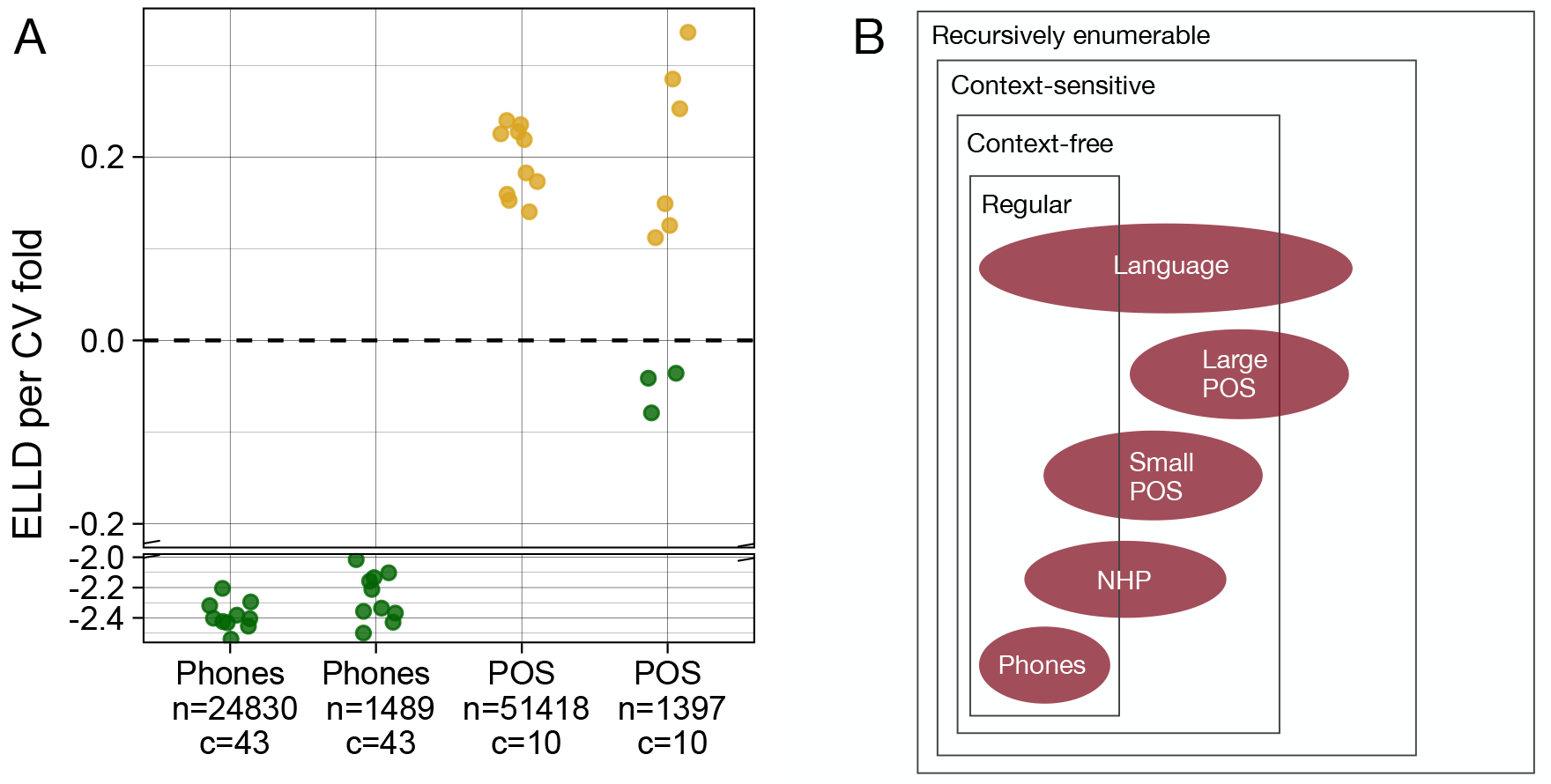
Human data and the Hierarchy of Grammars. **A**: Phone sequences are most likely generated by regular (pREG), parts-of-speech (POS) sequences most likely by supra-regular (pCFG) grammars, but when down-sampled to a similar number as in the nonhuman primate data (Figure 1A) parts of the POS sequences are most likely generated by a pREG. **B**: Most plausible position of primate communication systems in the inclusion hierarchy of grammar complexity, the Chomsky Hierarchy (Box 1, NHP: nonhuman primates).

## Discussion

The gap between human and animal intelligence may be vast in magnitude, yet decades of research continue to suggest it to be a difference in degree, not kind (*26*). The same argument has been made for complex language, with precursors described in the communication systems of nonhuman primates and other groups of animals. Syntax, however, has been the one main exception to the rule (*1*), mainly because of the apparent absence of specific kinds of hierarchical structures that permeate human language. So far it could not be shown that animals generate signal sequences that contain hierarchical, nested structures with arbitrary branching, as in the simple expression “They discussed biology curricula”.

Through developing and deploying a novel analysis tool to distinguish regular from context-free sequences, we show for the first time that animals generate call sequences that can be captured by supra-regular recursive grammars and hence share more similarities with human language than previously thought. Here, the key innovation has been to no longer analyze sequences as results of discrete mechanisms but as products of probabilistic processes. We deployed this tool to trawl through large corpora of primate call sequences, containing two hominid species (humans, bonobos) and two monkey species (olive colobus, marmosets). Across all four species we found parts of the sequences to be most likely generated by probabilistic grammars of supra-regular capacity. This narrows the gap between animal and human communication among primates. We furthermore found that this gap might be even narrower once substantially larger animal call corpora are considered.

In line with what has been observed with discrete grammars, both human and animal data straddle different complexity sets (Figure 2B). For human language this is generally assumed to be caused by fundamental differences between the type of rules that combine sounds and the type of rules that combine words (or parts of speech), but from all we know, no such distinction is operative in nonhuman primates (*27*). In light of this, a more likely explanation for the fact that nonhuman primate straddle complexity sets is that supra-regular mechanisms, while available, are only partially exploited in call sequences — perhaps only in certain contexts or because of learning constraints. Similar forces might also drive the distribution of human language over distinct complexity sets, above and beyond the differences between sound and word sequences. For example, speakers tend to under-exploit supra-regular recursion in child-directed speech (*28, 29*), i.e. speech tailored to facilitate language learning by children. Similarly, examining the evolutionary dynamics of complex sentences (sentences with at least two clauses such as “When I joined the meeting, they discussed biology curricula”), a recent study found that the grammars of very diverse languages systematically under-exploit supra-regular mechanisms, favoring flat concatenations instead (*30*). Since full exploitation is clearly attested as well, this would explain why parts of linguistic data fall into regular and other parts into supra-regular domains.

One potential limitation of our findings is that they might depend on how call sequences in primate species are discretized, i.e. what inventory is assumed for a species. Whilst, in the case of the olive colobus monkeys, the overall pattern is the same under two radically different discretizations (*c* = 2 vs. *c* = 9 calls; *Supplementary Figure S1*), more validation is needed against different discretizations in the other species as well.

An additional limitation is that we cannot assess what the precise resulting hierarchical structures, such as those exemplified for bonobos in Figure 1B, correspond to. Perhaps the call groupings — e.g., in the bonobo structure [[*wi sc*] *ch*] — are similar to word groupings in humans and reflect how meanings are composed (as in [[*cell biology*] *curricula*]). Recent evidence for widespread meaning compositionality in bonobos (*18*) makes this plausible, although much experimental work is needed to elucidate how exactly formal hierarchy maps to meaning composition. Alternatively, the groupings might reflect frequent associations that arise in response to similar contexts or when retrieving similarly-sounding calls for production. Yet another possibility is that the groupings reflect a purely formal structure, similar to the syntax of music in humans or song in animals. Disentangling these possibilities requires much further research. In the case of language, distinguishing meaning and structure is nontrivial either and progress mostly relies on neuro-imaging (*31, 32*) at a level that is not within immediate reach for research with nonhuman primates.

Despite these limitations, it is clear that the traditional view needs to be revised: some parts of call sequences of nonhuman primates are in fact most likely generated by supra-regular grammars. The absence of a supra-regular capacity has long been held to be the single-most important demarcation line in hominin evolution. Work with artificial visual stimuli challenged this (*11–13*), but it remained unclear whether the experiments merely showed that animals can be trained in a supra-regularity capacity or whether this taps into a naturally occurring behavior. Our present work addresses this longstanding conundrum, showing that the supra-regular capacity is indeed put to use in communication, at least for some parts of the call sequences of three primate species.

Our findings reset the agenda for research on the evolution of language: if not syntax, what is the critical step that enabled the evolution of language from a primate-like communication system? One candidate is the striking difference in the innovation of individual words. Humans constantly change the form and meaning of words, whereas the form and meaning of calls seem relatively fixed. This ultimately allows us to apply a finite lexicon to a rapidly changing world (*33–36*), for example, adopting the word for a drone (a male bee) to refer to an unmanned aerial vehicle. This flexibility does not rely on syntax, but on more fundamental mechanisms of human creativity. Our findings suggest that in order to unravel the evolutionary origins of language, it is not enough to focus on syntactic complexity as a demarcation line and it is necessary to pay attention to other aspects of human language such as communicative innovation.

## Author contributions

R.N. and B.B. designed research and experimental setup. R.N., Q.G., A.B, J.B., M.B., S.T., M.S., and K.Z. curated data. R.N. performed the investigation, experiments, and wrote relevant code. C.C., L.S.K. advised on methodology. L.S.K. advised on mathematical soundness. R.N., B.B., and K.Z. wrote the paper. R.N., B.B., S.T., C.C., L.S.K. and K.Z. edited the paper. B.B. and K.Z. supervised the work. All authors read and reviewed the paper.

## Data and materials availability

We provide all necessary code and data to replicate our experiment in the following OSF storage *link available on publication*. We further provide the codebase by itself for researchers to use for their own experiments in the following repository: *link available on publication*.

## Supplementary materials

Materials and Methods

Supplementary Text S1 to S3

Figs. S1 to S3

Table S1

References *(36-44)*

## Supplementary Materials

### Materials and Methods

#### Datasets

For the primate datasets, call annotations were done by the data providers (Table S1). Sequences that contained symbols which occurred only once in the dataset were removed during pre-processing. The human parts-of-speech (POS) data were taken from the British National Corpus (*37*). This corpus consists of a mix of written text and spoken conversations of British English. POS describes a linguistic class of word category, such as “noun”, “verb”, etc. We used the coarse-grained list of POS tags found in the British National Corpus, consisting of 9 categories. We took the POS-level annotation as provided in the corpus and used the “PUNCT” tag in order to split the utterances in the corpus into manageable sequences.

For the phonological data, we modeled single words as sequences and individual phones as symbols. A phonological annotation in the Buckeye Corpus for “Columbus” became the sequence “k ow l ah m b ah s”, where whitespace indicates the separation of symbols.

**Table S1.** Datasets used.

| Animal | Sequences | Provenance |
| --- | --- | --- |
| Common Marmoset (*Callithrix jacchus*) | 5800 | A. Bosshard, J. Burkart, (16) |
| Olive Colobus Monkey (*Procolobus verus*) | 1633 | Quentin Gallot |
| Bonobo (*Pan paniscus*) | 1107 | M. Berthet, S. Townsend, M. Surbeck |
| Human (English parts-of-speech) | 51’418 | BNC (37) |
| Human (English Phonology) | 24’839 | Buckeye Corpus (38) |

#### Methods

We first induced pREGs and pCFGs in an unsupervised manner. For pREGs we trained *n*-gram models of orders *k* = 1 … 6 on the training data split for each dataset and fold. We found that training orders of *k* > 6 did not yield any improvements in log-likelihoods of held-out data for any of the datasets. To account for unseen tokens we applied modified Kneser–Ney smoothing (*39, 40*) through the KenLM Python library (*41, 42*) for *k* > 1 (*Supplementary Text S1*).

For pCFGs, we adapted existing code (*43*) that induces an unbounded fully connected grammar (*Supplementary Text S2*) with a Bayesian inference model (*44*) (*Supplementary Text S1*). To allow efficient algorithmic parsing we represented each grammar in Chomsky Normal Form. Since the search space for pCFGs is technically infinite, we manually swept over the number of non-terminals *k*, finding highest log-likelihoods on held-out data in cross-validation for *k* = 6 … 20 in the nonhuman primate datasets and *k* = 6 … 35 in the human dataset. We did not restrict *k* to 6 … 35 a priori; models with *k* > 35 or *k* < 6 simply performed worse. In addition, we perform sensitivity analyses on the Dirichlet prior used in the induction model (*44*), finding that concentration parameters *β* = .07 … .2 generate the highest log-likelihoods on held-out data in cross-validation but specific values in this range make no appreciable difference. Again, as in the case of *k*, we searched for *β* individually for each dataset.

During training, we generated log-likelihoods over grammar parameters for each epoch. We monitored conversion by splitting the held-out testing data into two testing subsets, *test*_1_ and *test*_2_, with no overlap between the two sets. Most models converged after around 300–400 epochs. We stored the log-likelihood of *test*_2_ at the epoch with the highest log-likelihood for *test*_1_ and vice versa. This was only done to monitor conversion. For results we take the log-likelihood of the model at the last epoch, irrespective of the stored log-likelihoods of earlier epochs. The models for our primate datasets were trained for 600 epochs, while the models for our human datasets were trained for 500 epochs (to save on computational resources).

For each dataset we report the *expected log-likelihood difference* (ELLD) per *k*-fold. We derived this by performing a *k*-fold cross-validation split (*k*=20 for non-human primate data, *k*=10 for all others) for each dataset. Each training set was then used to induce a pCFG and a pREG model and we maximized the likelihood for each based on the *training data*. We next derived the log-likelihood of each model on the unseen data by taking the sum of their sequence log-likelihoods on the *held-out data* of the specific *k*-fold they were generated with. This defined a log-likelihood for each model type for each *k*-fold, *based on unseen data*. We then calculated the difference between these log-likelihoods (equivalent to a likelihood ratio) under the best pCFG and the best pREG for each *k*-fold and then took the mean of this difference. In other words, for each sequence *s*_*i*_ in each fold, we maximized the log-likelihood for the probability vectors **p** and sizes *k* (*n*-gram size or number of non-terminals) of each grammar *G* and then computed the expected difference over all sequences:

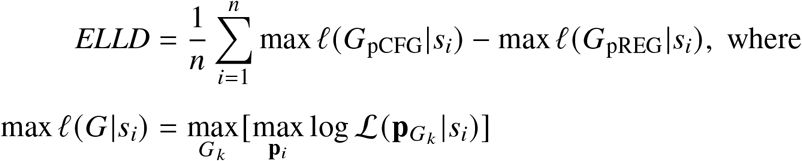

We use maximum likelihood for pREGs and maximum a posteriori (MAP) for the pCFGs (*43, 44*).

### Supplementary Text S1: Estimating Sequence Probabilities

Sequence probabilities for *n*-gram models (pREG) are calculated as the product of all symbol probabilities including the end-of-sequence symbol. For the *n* = 1 *n*-gram model this simply comes to 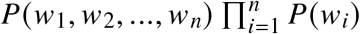 whereas for *n*-gram models where *n* > 1 this comes to:

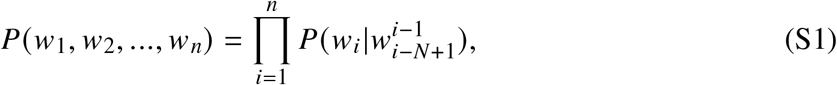

where *w*_*n*_ is the end-of-sequence symbol, *N* is the order of *n*-gram, and the context for *w*_1_ is the start-of-sequence symbol.

Our *n*-gram models are Kneser–Ney smoothed (*40, 41*). In essence, Kneser–Ney smoothed *n*- gram models consider counts of lower orders of *n* if the current context has not been observed. This makes the model more reliable. However, they are still fundamentally *n*-gram models and limited to a fixed linear context.

For pCFGs, the probability of a sequence *P*(*s*) is calculated as the sum of probabilities of all possible parses *P*(*t*) for the sequence:

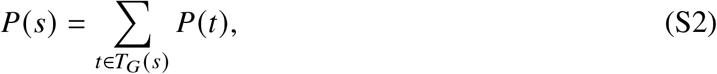

while the probability of a parse *P*(*t*) is defined as the product of the probabilities of all rule applications used to produce the parse.

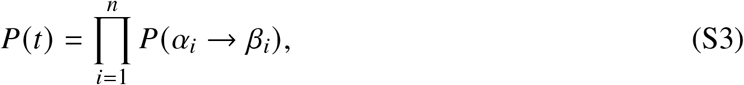

where *P*(*α*_*i*_ → *β*_*i*_) is the conditional probability of choosing rule *α*_*i*_ → *β*_*i*_ given *α*_*i*_ and, assuming that the parse is a possible parse of the grammar G: *t* ∈ *T*_*G*_ and that the parse contains rules: *α*_1_ → *β*_1_, *α*_2_ → *β*_2_, …., *α*_*i*_ → *β*_*i*_.

### Supplementary Text S2: A Probability Function for Sequences generated by pCFGs

We enforce the probability function of the pCFG to be distinct from that of our pREG model. This is done by assuming an unbounded fully-connected pCFG in Chomsky Normal Form (CNF) with no empty emissions (*23*). Under an unbounded fully-connected pCFG in CNF any binary combination of non-terminals is reachable from any given non-terminal and any terminal node may be emitted by any non-terminal node. Hence, any of the above rules have a non-zero probability. Finally, there are no restrictions on recursion depth (hence unbounded). The crucial factor is the fully-connected property and lack of empty emissions and recursion depth. The usage of Chomsky Normal Form is an engineering convenience and not strictly necessary.

This enforces a probability function which cannot be captured by a probabilistic regular grammar, such as an *n*-gram model (*23*). In fact, the inverse is also the case. A fully connected pCFG cannot capture the probability distribution over sequences that is generated by an *n*-gram model (pREG).

Probabilistic grammars model a probability distribution over possible sequences of symbols. We call this a *probabilistic language*. Under no restrictions, the *probabilistic languages* modeled by *n*-gram models (pREGs) are a subset of those modeled by unrestricted pCFGs. However, the *probabilistic languages* modeled by fully-connected pCFGs are *disjoint* from those modeled by *n*-gram models (*23*). The two classes of models allow the same sequences, but they can never converge on the same probability distributions over those sequences.

We use this disjoint property to our advantage, as it allows us to leverage likelihood as a measure. A model that is closer to the actual probability distribution that generated the data will yield the higher likelihood over aggregated data.

Enforcing a fully-connected pCFG is a diagnostic tool to distinguish between a pCFG and a pREG and should not be confused with an assumption about the underlying grammar. The underlying context-free mechanisms that are detected may not be fully connected in the strict sense and still be disjoint from the set of pREGs. However, to our knowledge the fully connected pCFG is the only diagnostic tool that can be shown to be disjoint from pREG under all circumstances.

### Supplementary Text S3: Validation of the likelihood method

We used sequence data generated from artificial pREGs (*k* = 2 … 5) and pCFGs (*k* ∈ *{*6, 9, 12, 15*}*)) to validate our methodology. For each value of *k*, we generated a grammar with a symbol vocabulary of size *c*, (*c* ∈ *{*6, 9, 12, 16*}*). For each set of parameters, we generate three distinct probability distributions *P*_*i*_. From each of these probability distributions, we generated datasets of *n* sequences where *n* ∈ *{*6000, 3000, 1500*}* for each grammar type (pREG and pCFG). This leaves us with 144 datasets for each grammar type (|*k* | × |*c*| × |*P*| × 3, where |*k* | = 4, |*c*| = 4, |*P*| = 3). We then performed our standard experiment as outlined in *Materials and Methods* with a *k*-fold split of *k* = 10. With these settings, we generated 10 ELLD values for each dataset. Due to the large number of trials, we do do not optimize the *k* parameter and use a single Dirichlet (*β* = 0.1) prior for the pCFG models in these experiments. The results of these experiments are plotted in Fig. S3, with all scripts and generated data available at [OSF URL].

### Supplementary Figures

**Figure S1.**
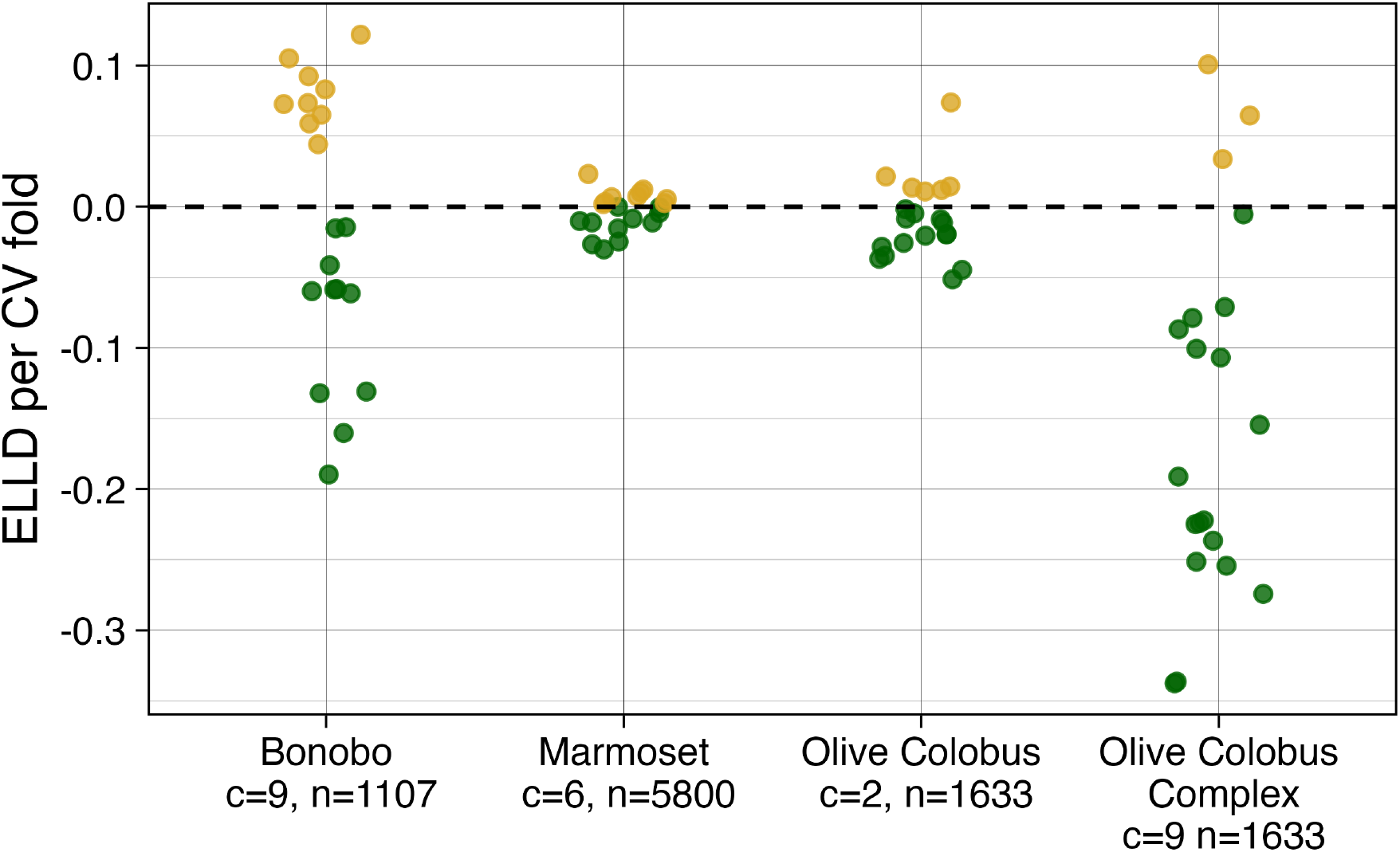
Results are stable across different symbol inventories. Even under a different vocabulary discretization that splits up the two Olive Colobus calls into nine distinct sub-calls, based on acoustic variation, there are still parts of the data that are better modeled by pCFGs.

**Figure S2.**
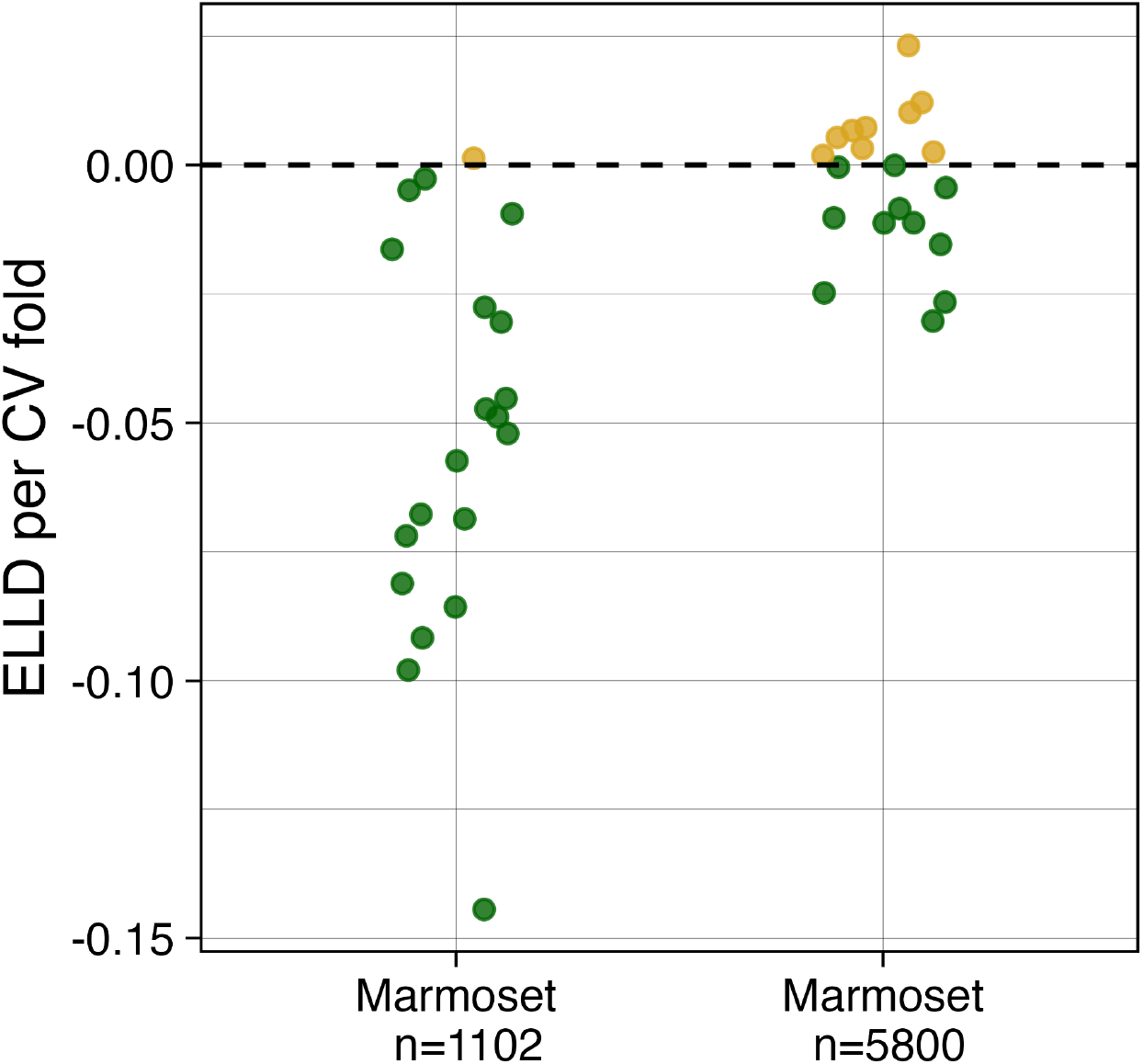
Approach is sensitive to data-size. A comparison between the Marmoset data and a down-sampled corpus of the same data shows that smaller data-sets skew the result towards pREG.

**Figure S3.**
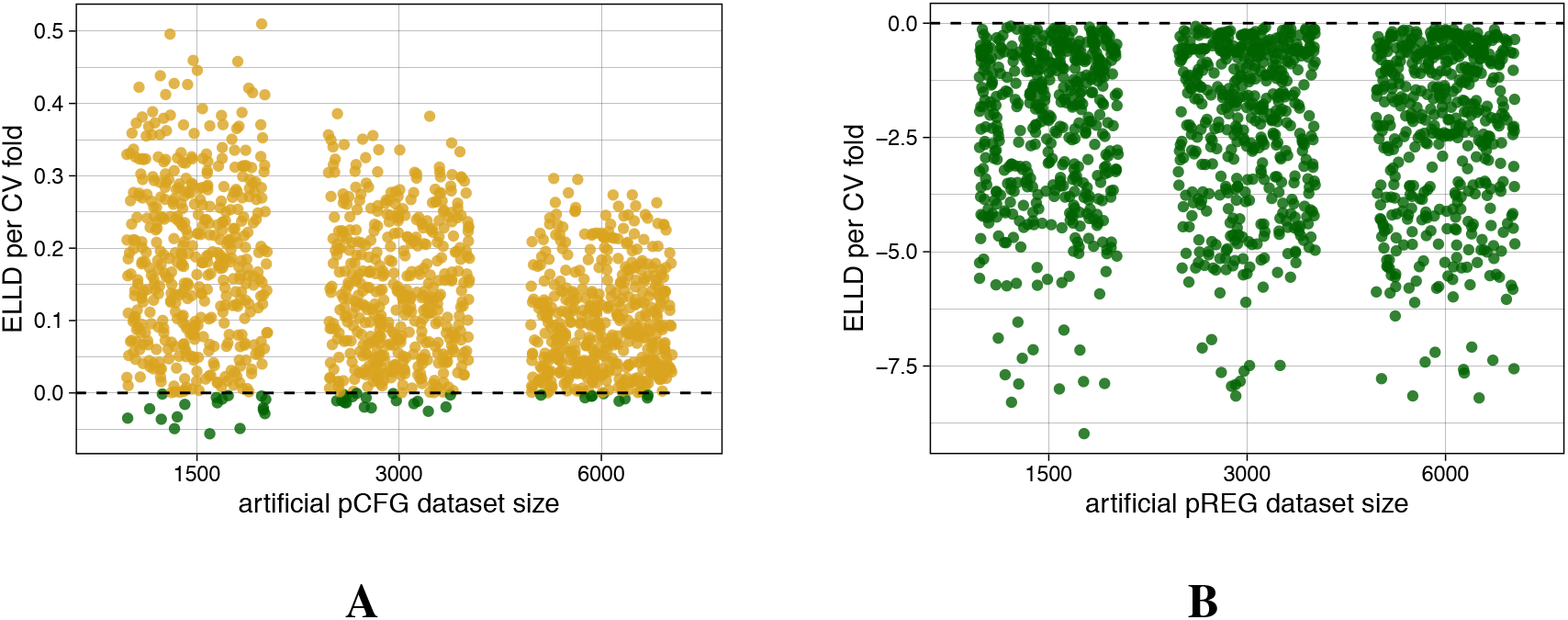
Validation Experiments. The outcome of 288 validation experiments (144 per class of grammar; *Supplementary Text S3*) show that the proportion of cross-validation folds better modeled by pCFGs increases with the number of sequences for the pCFG condition (**A**), but not for the pREG condition (**B**).

## References and Notes

1. M. D. Hauser, N. Chomsky, W. T. Fitch, The Faculty of Language: What It Is, Who Has It, and How Did It Evolve? Science (New York, N.Y.) 298, 1569–1579 (2002).

2. W. T. Fitch, Toward a Computational Framework for Cognitive Biology: Unifying Approaches from Cognitive Neuroscience and Comparative Cognition. Physics of Life Reviews 11, 329–364 (2014).

3. M. B. Everaert, M. A. Huybregts, N. Chomsky, R. C. Berwick, J. J. Bolhuis, Structures, Not Strings: Linguistics as Part of the Cognitive Sciences. Trends in Cognitive Sciences 19, 729–743 (2015).

4. A. D. Friederici, N. Chomsky, R. C. Berwick, A. Moro, J. J. Bolhuis, Language, Mind and Brain. Nature Human Behaviour 1 (10), 713–722 (2017), doi:10.1038/s41562-017-0184-4.

5. A. D. Friederici, Evolutionary Neuroanatomical Expansion of Broca’s Region Serving a Human-Specific Function. Trends in Neurosciences 46 (10) (2023), doi: 10.1016/j.tins.2023.07.004.

6. N. Chomsky, Three models for the description of language. IRE Transactions on Information Theory 2 (3), 113–124 (1956), doi:10.1109/TIT.1956.1056813.

7. N. Chomsky, On certain formal properties of grammars. Information and Control 2 (2), 137–167 (1959), doi:10.1016/S0019-9958(59)90362-6.

8. N. Chomsky, M. Schützenberger, The Algebraic Theory of Context-Free Languages*, in Computer Programming and Formal Systems, P. Braffort, D. Hirschberg, Eds. (Elsevier), vol. 35 of Studies in Logic and the Foundations of Mathematics, pp. 118–161 (1963), doi:10.1016/S0049-237X(08)72023-8.

9. S. Müller, Grammatical theory, no. 1 in Textbooks in Language Sciences (Language Science Press, Berlin) (2023), doi:10.5281/zenodo.7376662.

10. T. Graf, Subregular linguistics: bridging theoretical linguistics and formal grammar. Theoretical Linguistics 48 (3-4), 145–184 (2022), doi:10.1515/tl-2022-2037.

11. S. Ferrigno, S. J. Cheyette, S. T. Piantadosi, J. F. Cantlon, Recursive sequence generation in monkeys, children, U.S. adults, and native Amazonians. Science Advances 6 (2020), doi: 10.1126/sciadv.aaz1002.

12. R. Malassis, S. Dehaene, J. Fagot, Baboons (Papio papio) Process a Context-Free but Not a Context-Sensitive Grammar. Scientific Reports 10 (2020), doi:10.1038/s41598-020-64244-5.

13. X. Jiang, et al., Production of Supra-regular Spatial Sequences by Macaque Monkeys. Current biology : CB 28 12, 1851–1859.e4 (2018), doi:10.1016/j.cub.2018.04.047.

14. C. Girard-Buttoz, et al., Chimpanzees Produce Diverse Vocal Sequences with Ordered and Recombinatorial Properties. Communications Biology 5 (1), 410 (2022), doi:10.1038/s42003-022-03350-8.

15. Q. Gallot, C. Depriester, S. Moran, K. Zuberbühler, A Non-Human Primate Combinatorial System for Long-Distance Communication. iScience 27 (11), 111172 (2024), doi: 10.1016/j.isci.2024.111172.

16. A. B. Bosshard, et al., Beyond Bigrams: Call Sequencing in the Common Marmoset (Callithrix Jacchus) Vocal System. Royal Society Open Science 11 (11), 240218 (2024), doi: 10.1098/rsos.240218.

17. M. Leroux, et al., Call Combinations and Compositional Processing in Wild Chimpanzees. Nature Communications 14 (1), 2225 (2023), doi:10.1038/s41467-023-37816-y.

18. M. Berthet, M. Surbeck, S. W. Townsend, Extensive Compositionality in the Vocal System of Bonobos. Science 388 (6742), 104–108 (2025), doi:10.1126/science.adv1170.

19. S. M. Shieber, Evidence Against the Context-Freeness of Natural Language. Linguistics and Philosophy 8, 333–343 (1985), doi:10.1007/BF00630917.

20. G. Jäger, J. Rogers, Formal Language Theory: Refining the Chomsky Hierarchy. Philosophical Transactions of the Royal Society B: Biological Sciences 367, 1956–1970 (2012).

21. J. E. Hopcroft, J. D. Ullman, Introduction to Automata Theory, Languages, and Computation (Addison-Wesley) (1979).

22. S. Greibach, A note on undecidable properties of formal languages. Mathematical systems theory 2, 1–6 (1968), doi:10.1007/BF01691341.

23. L. S. Krapp, R. Nitschke, Contributions to the hierarchy of probabilistic languages (2026), arxiv preprint.

24. J. Heinz, W. Idsardi, What Complexity Differences Reveal About Domains in Language*. Topics in Cognitive Science 5 (1), 111–131 (2013), doi:10.1111/tops.12000.

25. J. Chandlee, J. Heinz, Computational Phonology, in Oxford Research Encyclopedia of Linguistics, M. Aronoff, Ed. (Oxford University Press) (2017), doi: 10.1093/acrefore/9780199384655.013.116.

26. C. Darwin, COMPARISON OF THE MENTAL POWERS OF MAN AND THE LOWER ANIMAL—continued, Cambridge Library Collection - Darwin, Evolution and Genetics (Cambridge University Press), p. 70–106 (1871/2009).

27. K. Collier, B. Bickel, C. P. van Schaik, M. B. Manser, S. W. Townsend, Language Evolution: Syntax before Phonology? Proceedings of the Royal Society B: Biological Sciences 281 (1788) (2014), doi:10.1098/rspb.2014.0263.

28. A. Perfors, J. B. Tenenbaum, E. Gibson, T. Regier, How recursive is language? A Bayesian exploration, in Recursion and Human Language, H. van der Hulst, Ed. (De Gruyter Mouton, Berlin, New York), pp. 159–176 (2010), doi:10.1515/9783110219258.159.

29. A. Perfors, J. B. Tenenbaum, T. Regier, The learnability of abstract syntactic principles. Cognition 118, 306–338 (2011), doi:10.1016/j.cognition.2010.11.001.

30. C. Y. Meloni, et al., Linguistic evolution minimizes the amount of syntactic hierarchy in grammar. bioRxiv pp. 2025–07 (2025).

31. L. Pylkkänen, The Neural Basis of Combinatory Syntax and Semantics. Science 366 (6461), 62–66 (2019), doi:10.1126/science.aax0050.

32. R. Riveland, A. Pouget, L. Driscoll, The Compositionality Continuum as a Principle for Studying the Neural Basis of Intelligence. Nature Neuroscience pp. 1–14 (2026), doi:10.1038/s41593-026-02382-1.

33. W. von Humboldt, Ü ber Die Verschiedenheit Des Menschlichen Sprachbaus Und Ihren Einfluss Auf Die Geistige Entwickelung Des Menschengeschlechtes (Dümmler, Berlin) (1836).

34. I. Meir, Topic-Open-Endedness: Why Recursion Is Overrated. Sign Language & Linguistics 23 (1-2), 258–271 (2020), doi:10.1075/sll.00051.mei.

35. B. Bickel, A.-L. Giraud, K. Zuberbühler, C. P. Van Schaik, Language Follows a Distinct Mode of Extra-Genomic Evolution. Physics of Life Reviews 50, 211–225 (2024), doi: 10.1016/j.plrev.2024.08.003.

36. B. Bickel, A.-L. Giraud, K. Zuberbühler, C. Van Schaik, Languages Evolve Ergodically: Clarifications and Responses. Physics of Life Reviews 57, 160–163 (2026), doi: 10.1016/j.plrev.2026.03.006.

37. BNC Consortium, The British National Corpus, XML Edition (Oxford Text Archive) (2007), https://www.natcorp.ox.ac.uk/.

38. M. Pitt, et al., Buckeye Corpus of Conversational Speech (2nd release) (Department of Psychology, Ohio State University) (2007), https://buckeyecorpus.osu.edu/.

39. R. Kneser, H. Ney, Improved backing-off for M-gram language modeling, in 1995 International Conference on Acoustics, Speech, and Signal Processing, vol. 1 (1995), pp. 181–184, doi: 10.1109/ICASSP.1995.479394.

40. S. F. Chen, J. Goodman, An empirical study of smoothing techniques for language modeling. Computer Speech & Language 13 (4), 359–394 (1999), doi: 10.1006/csla.1999.0128.

41. K. Heafield, KenLM: Faster and Smaller Language Model Queries, in Proceedings of the Sixth Workshop on Statistical Machine Translation (Association for Computational Linguistics, Edinburgh, Scotland) (2011), pp. 187–197, https://www.aclweb.org/anthology/W11-2123.

42. K. Heafield, I. Pouzyrevsky, J. H. Clark, P. Koehn, Scalable Modified Kneser-Ney Language Model Estimation, in Proceedings of the 51st Annual Meeting of the Association for Computational Linguistics (Volume 2: Short Papers) (Association for Computational Linguistics, Sofia, Bulgaria) (2013), pp. 690–696, https://www.aclweb.org/anthology/P13-2121.

43. L. Jin, F. Doshi-Velez, T. Miller, W. Schuler, L. Schwartz, Depth-bounding is effective: Improvements and evaluation of unsupervised PCFG induction, in Proceedings of the 2018 Conference on Empirical Methods in Natural Language Processing, E. Riloff, D. Chiang, J. Hockenmaier, J. Tsujii, Eds. (Association for Computational Linguistics, Brussels, Belgium) (2018), pp. 2721–2731, doi:10.18653/v1/D18-1292.

44. M. Johnson, T. Griffiths, S. Goldwater, Bayesian Inference for PCFGs via Markov Chain Monte Carlo, in Human Language Technologies 2007: The Conference of the North American Chapter of the Association for Computational Linguistics; Proceedings of the Main Conference, C. Sidner, T. Schultz, M. Stone, C. Zhai, Eds. (Association for Computational Linguistics, Rochester, New York) (2007), pp. 139–146, https://aclanthology.org/N07-1018/.

